# An investigation of the effects of α and β-frequency neural entrainment using tACS on phase-aligned TMS-evoked corticospinal excitability

**DOI:** 10.1101/2024.12.03.626574

**Authors:** Aikaterini Gialopsou, Stephen R. Jackson

## Abstract

Deep brain stimulation [DBS] is an effective treatment for many brain disorders (e.g., Parkinson’s disease), has a favourable adverse effect profile, and can be particularly effective for individuals with treatment resistant symptoms. DBS is, however, inaccessible for most individuals, is extremely expensive, and is not considered suitable for children and adolescents. For these reasons, non-invasive alternatives to DBS, such as transcranial magnetic stimulation [TMS], are increasingly being sought to treat brain health conditions. Unfortunately, current TMS approaches exhibit large intra- and inter-subject variability in their efficacy, which limits their use clinically. One likely reason for this is that TMS is invariably delivered without reference to ongoing brain activity (i.e., open loop). We propose that the efficacy of stimulation might be improved, and the variability of its effects reduced, if stimulation could be synchronised with ongoing brain activity. To investigate this, we used transcranial alternating current stimulation (tACS) to induce entrainment of brain activity at two frequencies (α=10 Hz and β=20 Hz), and we delivered single pulse TMS that was temporally aligned with the phase of each tACS oscillation. To investigate the effects of tACS-phase-aligned TMS we measured motor-evoked potentials (MEPs). Our findings confirm that for α-tACS and β-tACS both corticospinal excitability and inter-trial variability varied as a function of tACS phase. Importantly, however, the tACS phase angle that produced maximum TMS-evoked excitability was different for α-tACS and β-tACS; coinciding with the negative peak (trough) for α-tACS and the positive peak (peak) for β-tACS. These findings confirm that aligning non-invasive brain stimulation to ongoing brain activity may increase the efficacy of TMS and reduce the variability of its effects. However, our results illustrate that the optimal phase of the tACS cycle at which to deliver TMS may vary for different tACS frequencies.

## Introduction

DBS is a surgical procedure in which implanted electrodes deliver high frequency electrical stimulation to a targeted brain area to modulate dysfunctional brain network activity. Importantly, the mode-of-action of DBS would appear to depend upon neural entrainment, as studies have shown that during DBS neural firing rates become strongly phase-locked to the DBS pulse trains [1].

DBS is typically used only with treatment-resistant adults, is very expensive, is not suitable for children or adolescents, and is restricted in its availability. For this reason, non-invasive brain stimulation alternatives to DBS are being sought as these may have the potential to widen access to safe and effective brain stimulation treatments for a broad range of neurological and psychiatric conditions.

Non-invasive brain stimulation approaches such as TMS are increasingly being used to treat brain health conditions such as depression [2]. However, current TMS approaches have a number of limitations that may limit their use clinically. First, they cannot be used to directly stimulate deep brain areas that may play role in brain disorders, although they may be effective indirectly. Second, the effects of TMS are often highly variable [3], exhibiting considerable intra-subject and inter-subject variability, which limits their use clinically.

One likely contributing factor for the variability observed for non-invasive brain stimulation approaches such as TMS, is that current approaches most often adopt an open-loop, one-size-fits-all, methodology in which individuals each receive identical stimulation, without reference to ongoing brain activity. However, recent evidence strongly indicates that the efficacy of TMS is increased by adopting a closed-loop control strategy, in which stimulation is time-locked to ongoing physiological events within the brain [4].

Cortical activity is characterized by neural oscillations that are thought to reflect synchronised neural firing. These oscillations can be measured in using electroencephalography (EEG) or magnetoencephalography (MEG) and reflect spontaneous fluctuations in brain state, that are characterised by differences in cortical activity. Specifically, alterations in cortical excitability are shown to be associated with the instantaneous phase of cortical oscillations [5]. This may indicate a functional role for brain oscillations in organising patterns of neural firing in the brain. In the motor cortex, changes in power in α-band (8-13 Hz) and β -band (14-30 Hz) oscillations are associated with the preparation and initiation of voluntary movements [6-8].

Zrenner and colleagues used real-time, EEG-triggered, TMS to demonstrate a dependence between TMS-induced alterations in corticospinal excitability and the phase of the cortical α-rhythm [4]. They reported that larger MEP amplitudes were elicited when TMS was triggered to coincide with the negative peak (trough) of the α-rhythm, compared to when it was aligned with the positive peak [4]. See also [9,10]. Consistent with this finding, functional magnetic resonance imaging (fMRI) studies have demonstrated that sensorimotor activity (as measured by fMRI BOLD signal) is negatively correlated with the sensorimotor EEG α-rhythm [11].

Wischnewski and colleagues [12] also used real-time EEG-triggered TMS to investigate the relationship between TMS-induced increases in corticospinal excitability and the phase of the cortical α-rhythm or β-rhythm [12]. They targeted four phases (0^°^, 90 ^°^, 180 ^°^, and 270 ^°^) of the α and β rhythms with suprathreshold single-pulse TMS to the primary motor cortex. They reported phase-dependent modulation of TMS-induced corticospinal excitability for both the α-rhythm and β-rhythm. Importantly, they found that the relationship between EEG phase and TMS-induced corticospinal output differed for the α-rhythm or β -rhythm. Motor-evoked potentials (MEPs) were largest when TMS was aligned with the trough and rising phase of the α-rhythm compared to when TMS was aligned with either the peak or falling phase. By contrast, for the β-rhythm MEPs were largest when TMS was aligned with the peak and falling phase of the β-rhythm.

Studies such as that reported by Zrenner [4] and Wischnewski [12] demonstrate that closed loop control of non-invasive brain stimulation can be achieved by temporally aligning stimulation with the phase of brain oscillations measured and analysed in real-time using EEG. However, the real-time analysis of EEG and the accurate temporal prediction of instantaneous oscillatory phase is technically challenging. An alternative, and arguably simpler, method for delivering non-invasive brain stimulation in phase with neural activity may be through neural entrainment.

As noted above, the mode-of-action of DBS depends upon neural entrainment as during DBS neural firing become strongly phase-locked to the frequency of DBS pulse trains [1]. Importantly, entrainment of neural activity can also be achieved non-invasively using transcranial alternating current stimulation (tACS) [13]. tACS involves applying a weak sinusoidal electrical current between two or more electrodes attached on the scalp and is thought to induce entrainment of neural activity (in phase with the frequency of the tACS stimulation) and spike-timing dependent synaptic plasticity (STDP) leading to stimulation aftereffects [13]. Studies in non-human primate demonstrate that tACS can induce: (a) phase-entrainment of neural firing [14], (b) dose-dependent increases in neural bursting [15]; and (c) cell-class-specific entrainment of spiking activity [16]. Importantly, these studies indicate that while tACS induces phase-dependent entrainment of neural firing, it does not increase the neural spiking rate [14]. Studies in humans have demonstrated that β-frequency tACS can be used successfully to entrain cortical sensorimotor activity and as a result slow the execution of voluntary movement [17]. Furthermore, recent advances in the methods used to deliver tACS have shown that modulation of deep brain areas such as the hippocampus or striatum is possible using tACS [14] or a by using variant of tACS known as temporal interference stimulation (TIS) [18,19]. It is suggested that tACS might be used to normalise neural ‘noise’ in maladaptive brain networks linked to neurological disease [14] and open up the possibility of developing non-invasive alternatives to current surgical DBS procedures.

In the current study we used α- and β-frequency tACS to entrain cortical motor oscillations and delivered single pulses of above-threshold TMS to coincide with the peak (0^°^), falling phase (90^°^), trough (180^°^), or rising phase (270^°^) of the α or β phase cycle. TMS-evoked corticospinal excitability was measured using electromyography (EMG). We hypothesised that: (a) there would be phase-dependent differences in the magnitude of TMS-evoked MEPs; (b) there would be phase-dependent differences the inter-trial variability of MEP amplitudes; and (c) phase-dependent effects might be different for tACS entrainment at α and β frequencies.

## Methods

Participants were recruited to take part in two separate sessions that each involved the application of combined tACS and TMS over the primary motor cortex. In each case MEP amplitudes and trial-by-trial variability were the primary outcome measure. The order of the sessions was counterbalanced to control for potential sequence effects.

### Participants

Data were collected from 15 right-handed participants (10 female, mean age ± SD: 26.6 ± 5.7 years), all of whom had no history of neuropsychological disorders (Table 1). Prior to the study, each participant provided written informed consent and completed a TMS safety questionnaire either online or in person. The study protocol was reviewed and approved by an appropriate local ethical review committee (School of Psychology, University of Nottingham). All participants completed both sessions of the study, which were counterbalanced and scheduled to occur at least three days apart.

**Table 1.** Participant information including motor threshold (MT) values recorded for session one (S01) and session two (S02).

| ID | Age | Sex | MT % S01 | MT % S02 |
| --- | --- | --- | --- | --- |
| HP01 | 24 | F | 70 | 70 |
| HP02 | 27 | F | 67 | 72 |
| HP03 | 24 | F | 75 | 75 |
| HP04 | 34 | M | 76 | 76 |
| HP05 | 23 | F | 72 | 72 |
| HP06 | 33 | M | 72 | 72 |
| HP07 | 19 | F | 68 | 68 |
| HP08 | 28 | F | 70 | 70 |
| HP09 | 18 | F | 76 | 76 |
| HP10 | 37 | M | 74 | 74 |
| HP11 | 34 | F | 76 | 76 |
| HP12 | 28 | F | 56 | 56 |
| HP13 | 25 | M | 70 | 70 |
| HP14 | 22 | F | 56 | 56 |
| HP15 | 23 | M | 54 | 54 |

### Procedure

Participants were seated in a chair with their right arm and elbow placed in a comfortable position on the desk in front of them. EMG was recorded using disposable Ag-AgCl muscle electrodes (H124SG Covidien, 23.9 mm diameter). Electrodes were placed securely on the right hand in a belly-tendon montage targeting the first dorsal interosseus (FDI) muscle, with the ground electrode placed on the right ulna. EMG signals were amplified and bandpass filtered (10Hz-2kHz, with a sampling rate of 2kHz) and then digitized using Brain Amp ExG (Brain Products, GmbH, Gilching, Germany) controlled by Brain Vision Recorder (Brain Products, GmbH, Gilching, Germany).

TMS hotspotting, thresholding, and single-pulse stimulation were delivered using a MagStim Bistim^2^ stimulator (Magstim, Whiteland, Dyfed, UK) with a 70 mm figure-of-8 coil coupled to a BrainSight neuronavigation system (Rogue Research Inc., Montreal, Quebec, Canada). Neuronavigation was based on a template MRI scan (MNI-152) to allow for accurate coil orientation and location over the left M1. Before delivering pulses to the scalp, participants had the opportunity to ask questions and feel a test pulse of TMS on the palm of their hand once the procedure was explained to them. Once ready, suprathreshold pulses were delivered to the left M1 hand area ‘hotspot’, defined as the location that consistently evoked the largest MEPs in the right FDI muscle. Coil orientation was maintained at 45 degrees from the midline with current flowing in a posterior-anterior direction, which has been shown to evoke optimal MEP responses in the FDI muscle [20]. These pulses were recorded on BrainSight and once the hotspot was found, this area was landmarked to ensure this hotspot area was consistently targeted at the correct orientation throughout the experiment and allowed participants to take breaks if they requested one.

Once the hotspot was identified, this area was marked to ensure the correct placement of the tACS electrode. The tACS neurostimulator (NeuroConn, Germany) used tACS sponge electrodes of size of 5cm × 7cm. The tACS electrode was placed over the marked area (left M1) while the other electrode was placed over Pz (according to 10-10 system [21]). The electrode dimension was 5cm × 7cm. The impedance of the electrodes was always below 10 KΩ for all the participants, which minimizes the tactile sensation. This was achieved by using conductivity gel and saline. The tACS electrodes were placed on the participants head and stabilized there using rubber straps.

Following the hotspotting and the placement of tACS electrodes, motor thresholds (MT) were determined for each participant using EMG responses from the first dorsal interosseous (FDI) muscle. MT was defined as the TMS stimulation intensity (% maximum stimulator output, MSO) required to elicit a peak-to-peak MEP amplitude of 200-500 µV in approximately 5 out of 10 trials. The high MT was selected to ensure clear MEP visibility throughout the sessions.

In addition to the tACS electrodes, two Ag-AgCl electrodes (H124SG Covidien, 23.9 mm diameter) were placed over the M1 tACS electrode using rubber straps. These electrodes were utilized to record real-time tACS oscillations via the Brain Amp ExG system (Brain Products, GmbH, Gilching, Germany).

A custom MATLAB script was employed to trigger and synchronize tACS stimulation with single-pulse TMS delivery. Each session consisted of four trials, each lasting 5 minutes, with optional breaks available to participants between sessions. At the onset of each trial, tACS and TMS were simultaneously triggered. TMS pulses were delivered with an intertrial interval of 4 ± 1 s, under two conditions: TMS-pulse or no TMS-pulse, randomly assigned across different tACS phases.

### Analysis

EMG data for each participant and trial was inspected visually using custom in-house software developed in MATLAB. Trials were excluded from further analysis if pre-contraction of the FDI muscle, or noise levels, exceeded 50 μV during the 500 ms window preceding onset of MEP. The peak-to-peak amplitudes of MEPs were quantified within this software. A custom MATLAB script was used to calculate the alignment of the tACS phase to each participant’s MEPs. The MEPs and their corresponding tACS phases were then visually inspected and categorized into four phases: peak, trough, falling edge, and rising edge. Median MEP amplitudes were calculated for each tACS phase (peak, falling edge, trough, rising edge) for each participant. Inter-trial variability was also assessed by calculating the coefficient of variation (CV) for each tACS phase (peak, falling edge, trough, rising edge) and for each individual. The mean of individual median MEP values and CV values for all participants was used to identify differences across tACS phases and between stimulation frequencies. Effect sizes based on standardised mean differences (Cochrane’s D) were calculated along with t-tests to evaluate phase-related differences in mean MEP amplitudes and CV values. By convention a Cochrane D value of 0.5 is taken to reflect a medium sized effect.

## Results

### Mean number of TMS pulses delivered

Initial t-tests confirmed that were no statistically significant differences in the mean number of MEPs collected across the different tACS phases and stimulation frequencies (p > 0.05). Relevant mean number of TMS pulses delivered for the α-tACS session were peak = 30, trough = 39, falling edge = 32, rising edge = 29. And for the β-tACS session, peak = 34, trough = 30, falling edge = 25, and rising edge = 29.

### Motor threshold

The mean motor threshold (MT%) for session 1 was 68.8 (7.54) % and for session 2 it was 69.1 (7.56) %. These means did not differ statistically (p > 0.05).

### MEP amplitude

Mean standardised (z-score) median MEP amplitudes for α-tACS and β-tACS entrainment are presented in Figure 2 (upper panels). For α-tACS entrainment the mean (± SD) MEP amplitude was -0.40 (± 0.8) at the peak, -0.05 (± 0.8) at the falling edge, 0.72 (± 0.8) at the trough, and -0.28 (± 0.7) at the rising edge. Statistical analyses confirmed that when TMS was aligned with the α-tACS trough then MEP amplitudes were larger than when aligned with the peak, falling edge or rising edge (minimum: Cohen D = 0.53; p < 0.05).

**Figure 1:**
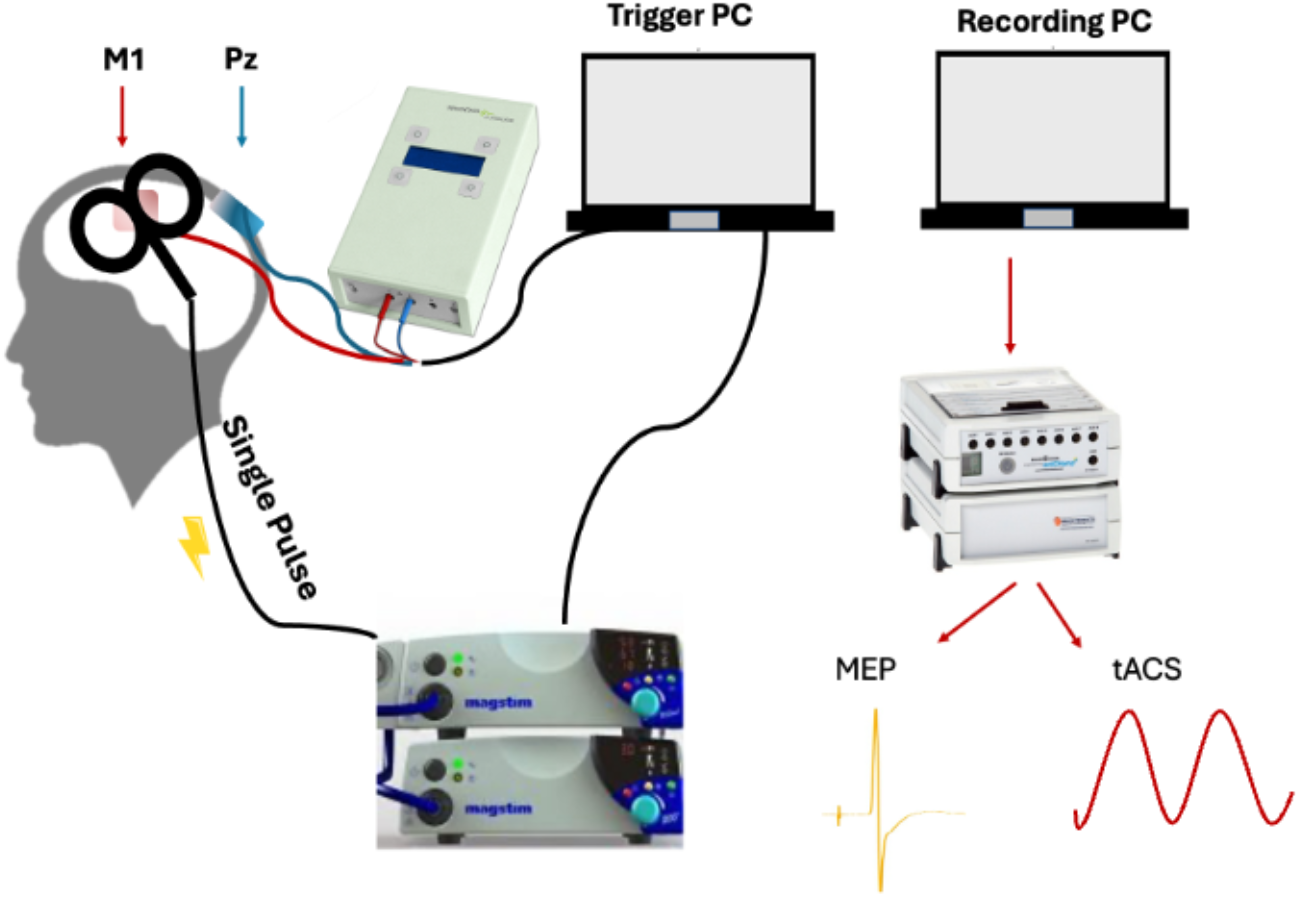
Experimental design. The tACS electrode was placed over the hand region of the left primary motor area M1 (as determine by TMS hotspotting) and the other tACS electrode was placed over the Pz. Electrodes were held in place by a rubber strap, with two EMG electrodes attached over the M1 tACS electrode. The TMS coil was placed over the M1. The trigger PC triggered and synchronized the tACS and TMS while the recording PC measured the MEPs from FDI and tACS oscillations from the M1 electrode.

**Figure 2:**
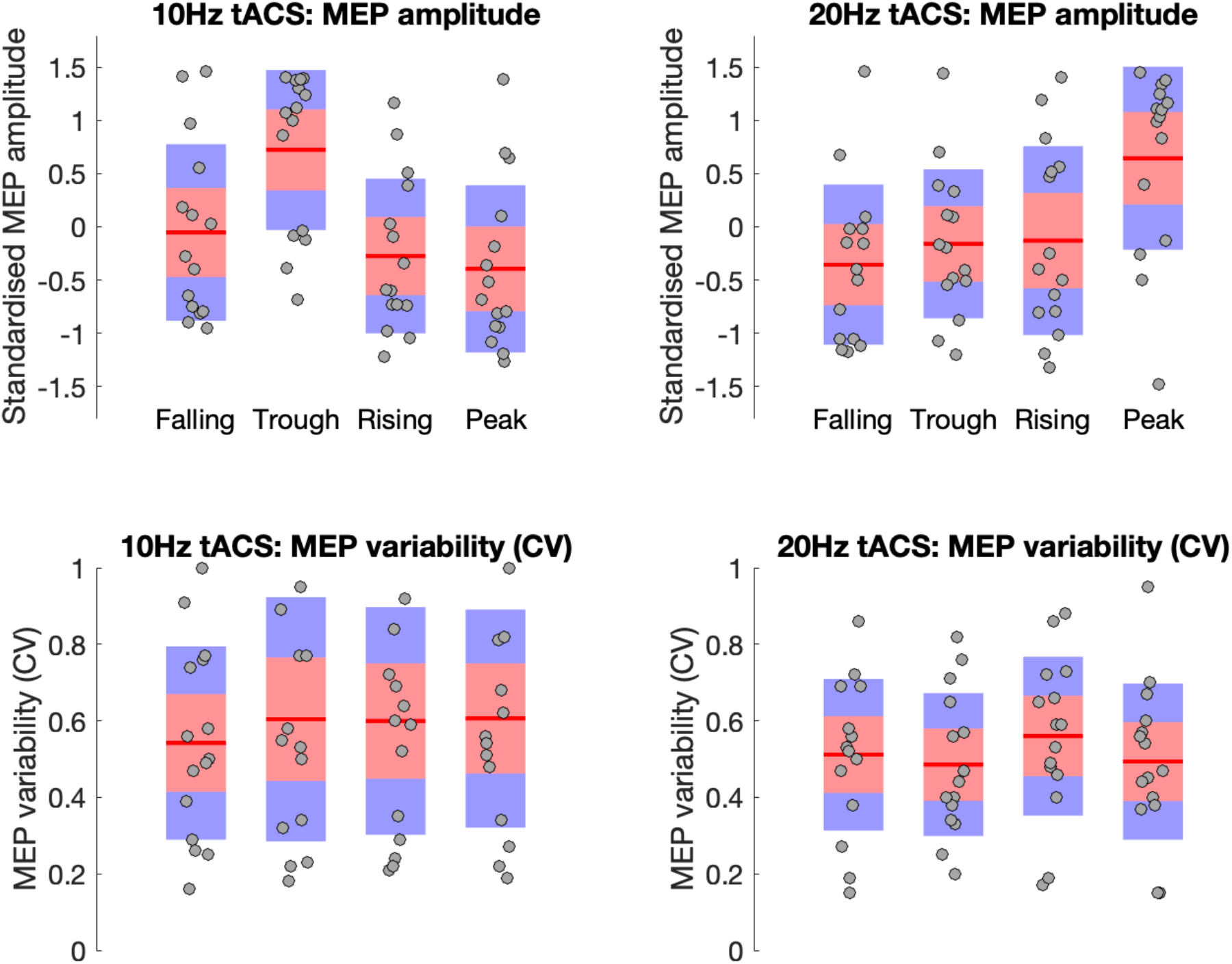
<u>Upper panel</u>: Boxplots showing the mean of standardised median MEP amplitudes during the α-tACS stimulation (left) and β-tACS stimulation (right). The grey dots indicate the individual mean of medians while the red lines indicate the median for each condition (Peak, Falling Edge, Trough, Rising edge). <u>Bottom panel</u>: Boxplots showing the mean coefficient of variation [CV] values during the α-tACS stimulation (left) and β-tACS stimulation (right).

For β-tACS entrainment the mean (± SD) MEP amplitude was 0.65 (± 0.9) at the peak, -0.36 (± 0.8) at the falling edge, -0.16 (± 0.7) at the trough, and -0.13 (± 0.9) at the rising edge. Statistical analyses confirmed that when TMS was aligned with the β-tACS peak then MEP amplitudes were larger than when aligned with the trough, falling edge or rising edge (minimum: Cohen D = 0.53; p < 0.05).

These findings confirm that TMS-induced corticospinal excitability varies as a function of tACS phase. Specifically, they confirm that MEP amplitude is maximal when TMS is aligned with the trough of the α-tACS phase cycle but with the peak when aligned with the β-tACS phase cycle. These results are consistent with previous results obtain ed using EEG-triggered TMS [4,12].

### MEP variability

Mean coefficient of variation (CV) values for α-tACS and β-tACS entrainment are presented in Figure 2 (lower panels). For α-tACS entrainment mean (± SD) CV values were 0.61 (± 0.3) at the peak, 0.54 (± 0.3) at the falling edge, 0.60 (± 0.3) at the trough, and 0.60 (± 0.3) at the rising edge. Statistical analyses confirmed that when TMS was aligned with the α-tACS falling edge the mean CV was reduced compared to when TMS was aligned to the α-tACS trough (D = -0.57, p < 0.05).

For β-tACS entrainment mean (± SD) CV values were 0.49 (± 0.2) at the peak, 0.51 (± 0.2) at the falling edge, 0.49 (± 0.2) at the trough, and 0.56 (± 0.2) at the rising edge. Statistical analyses confirmed that when TMS was aligned with the β-tACS trough the mean CV was reduced compared to when TMS was aligned to the β-tACS rising edge (D = -0.52, p = 0.06).

These findings confirm that trial-by-trial variability of TMS-induced increases in corticospinal excitability, as indexed by CV, varies as a function of both tACS phase and tACS frequency. Interestingly, these results indicate that the optimal target for delivering phase-aligned TMS may differ according to whether one aimed to maximise the efficacy of TMS stimulation (i.e., increase MEP amplitude) or minimise the variability of stimulation (i.e., decrease MEP CV). This can be seen in Figure 3. Inspection of this figure suggests that if one wanted to both increase efficacy of TMS and decrease variability, then one might align TMS with the falling edge of 10 Hz tACS rather than the trough.

**Figure 3:**
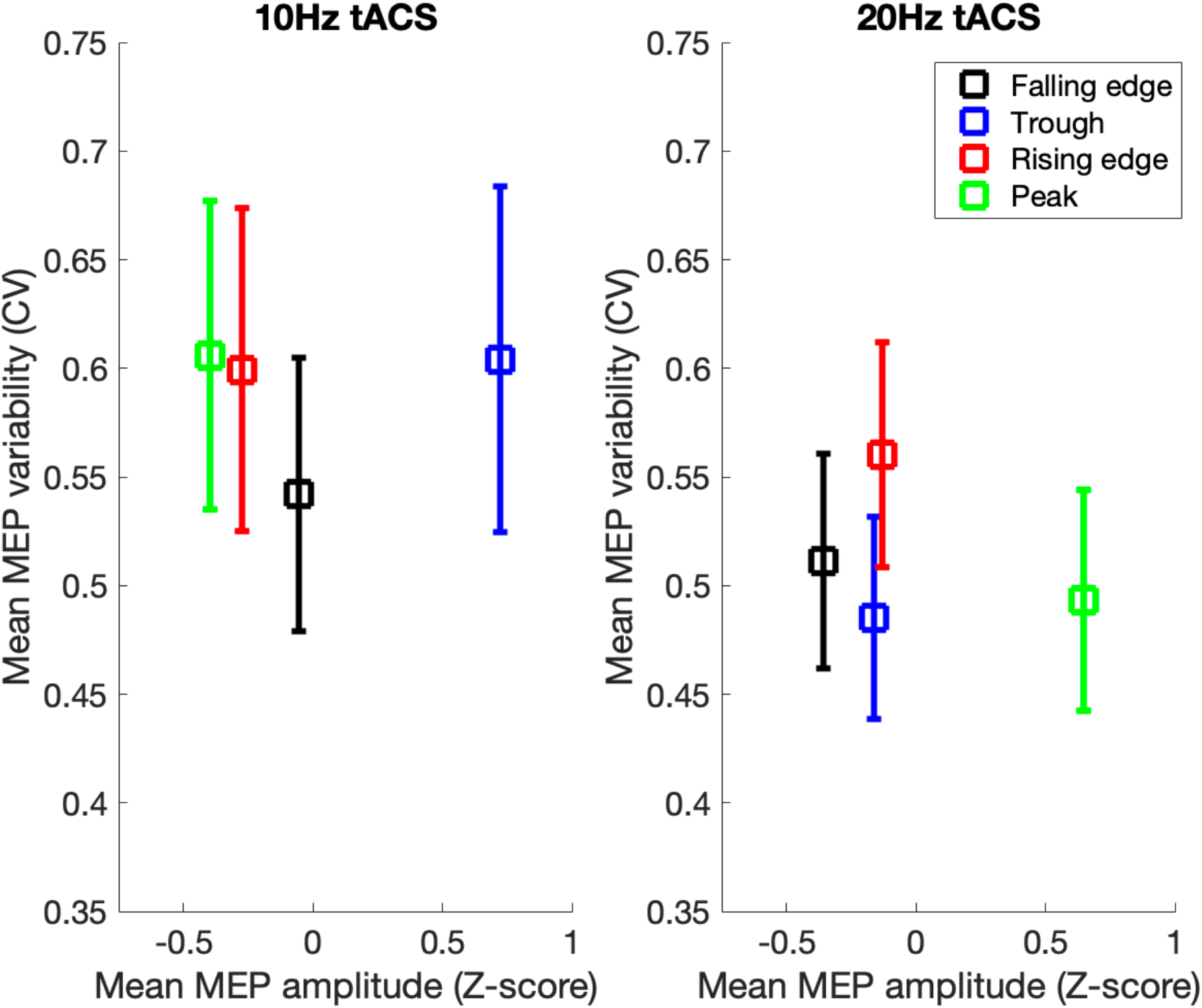
<u>Left panel</u>: shows the relationship between mean TMS-induced MEP amplitude (x-axis) and mean TMS-induced MEP variability when TMS was aligned to each phase of the α-tACS (10 Hz) phase cycle. Error bars are the standard error of the mean (sem). <u>Right panel</u>: shows the relationship between mean TMS-induced MEP amplitude (x-axis) and mean TMS-induced MEP variability when TMS was aligned to each phase of the β-tACS (20 Hz) phase cycle.

## Discussion

In this study we used α- and β-frequency tACS to entrain cortical motor oscillations and we delivered single pulses of TMS to induce increases in corticospinal excitability as measured by EMG motor-evoked potentials (MEPs). Importantly, on each trial the delivery of TMS was timed to coincide with the peak, falling phase, trough, or rising phase of the α or β phase cycle. Based on prior EEG-triggered TMS studies we hypothesised that there would be phase-dependent differences in the magnitude of TMS-evoked MEPs and that these phase-dependent effects would differ for α and β tACS entrainment [4,12]. Importantly, we also hypothesised there would be phase-dependent differences in the inter-trial variability of TMS-induced increases in MEP amplitude. Our results confirmed each of these hypotheses which are discussed in more detail below.

In this study we have assumed that tACS delivered at a given (target) frequency can be used to entrain brain activity at that target frequency in a phase-dependent manner. This assumption is supported by several converging lines of evidence. First, studies in non-human primates demonstrate phase-dependent alignment of neural spiking activity during tACS [14-16]. Note that in this case, the neural firing rate is not actually increased but instead, spiking is organised to occur in bursts that are aligned with the phase of the tACS stimulation [14]. Second, studies that have combined EEG recording with tACS stimulation support the proposal that tACS modulates EEG at the targeted frequency. Zaehle and colleagues used tACS delivered at participants’ individual α-frequencies, and they demonstrated that α-power was increased after tACS compared to after sham stimulation [22]. More recently, Helfrich and colleagues overcame the inherent technical difficulties of using EEG to concurrently measure the online effects of tACS and demonstrated that 10 Hz tACS enhances EEG amplitudes in the alpha band [23]. They also reported a narrowing of the spectral peak in the alpha band [23]. This may indicate that, consistent with the concept of tACS phase-aligned entrainment of brain oscillations, individual preferred α-frequencies of participants underwent a shift towards the target 10 Hz tACS frequency. Finally, studies that have used concurrent fMRI to investigate the effects of tACS also support the proposal that tACS entrains brain activity. Specifically, previous fMRI studies have shown that α-frequency power is negatively correlated with the fMRI BOLD signal [11,24,25]. For this reason, Vosskuhl and colleagues used fMRI to investigate the effects of α-tACS on the fMRI occipital BOLD response to visual stimulation [26]. They demonstrated that α-tACS reduced task-related BOLD activation compared to tACS-free periods [26]. Based on these converging lines of evidence, we feel confident in concluding that tACS can be used in our study to effectively entrain 10Hz and 20Hz brain oscillations.

As noted above, studies making use of real-time phase-aligned EEG-triggered TMS have demonstrated that TMS-induced increases in MEP amplitude are larger when TMS is triggered to coincide with the trough of the 10 Hz oscillation [4,12] but conversely, are larger when timed to coincide with the peak of the 20 Hz oscillation [12]. Consistent with these previous reports, we demonstrate here that TMS-induced increases in MEP amplitude are largest when timed to coincide with the trough of the α-tACS phase cycle or the peak of the β-tACS phase cycle. To the extent that brain oscillations may reflect functionally important fluctuations in brain state (e.g., changes in cortical excitability), our findings, together with those of the EEG-triggered TMS studies referred to above, highlight the impact that the frequency and phase alignment of ongoing brain oscillations may have on the efficacy of non-invasive brain stimulation.

We hypothesised that aligning TMS with the phase of tACS might not only increase the efficacy of TMS but would also increase the consistency of the effect, by reducing trial-by-trial variability. Our findings demonstrate that there were indeed phase-related differences in the trial-by-trial variability of TMS (as measured by CV) and that these differed for α-tACS and β-tACS stimulation. Specifically, we found that variability was reduced when TMS was aligned to the falling edge of the α-tACS phase cycle and to the trough of the β-tACS. This finding indicates that choosing precisely when to deliver non-invasive brain stimulation with respect to ongoing brain oscillations it may be important to consider not only the oscillation frequency and phase, but also whether the aim is to increase the efficacy of stimulation or reduce the variability of stimulation effects.

While this study provides evidence that tACS can be used to entrain corticospinal excitability in a frequency-specific and phase-specific manner, much still remains to understand with respect to the underlying mechanisms involved. tACS is generally thought to cause the dendrites and cell bodies of cortical neurons to alter their membrane potential towards depolarization or hyperpolarization in synchrony with the oscillation frequency of the stimulation. This suggests that tACS primarily acts to organise and synchronise neural firing rather than initiate firing. Consistent with this proposal, animal studies demonstrate that while spiking becomes synchronised with the phase of tACS, the overall rate of spiking does not increase [4]. Similarly, human fMRI studies show that while α-tACS modulates task-related BOLD activation compared to tACS-free periods, there was no effect of tACS independent of the visual task [26]. However, recent animal studies have demonstrated that tACS may lead to frequency-dependent cell-class-specific entrainment of spiking activity [[16]. This raises the possibility that entrainment of cortical activity might involve the selective activation modulatory local interneuron populations.

## Acknowledgements

Stephen Jackson is supported by research grants from MRC (T032588), EPSRC, Tourettes Action and Parkinson’s UK. Kat Gialopsou by research grants from MRC (T032588) and Parkinson’s UK. We would like to thank Patricia Radu for the help with the data collection during her PhD rotation.

